# PET studies of the glial cell marker TSPO in psychosis patients - a meta-analysis using individual participant data

**DOI:** 10.1101/228742

**Authors:** Pontus Plavén-Sigray, Granville J. Matheson, Karin Collste, Abhishekh H. Ashok, Jennifer M. Coughlin, Oliver D. Howes, Romina Mizrahi, Martin G. Pomper, Pablo Rusjan, Mattia Veronese, Yuchuan Wang, Simon Cervenka

**Affiliations:** Department of Clinical Neuroscience, Center for Psychiatry Research, Karolinska Institutet and Stockholm County Council, SE-171 76 Stockholm, Sweden.; IoPPN, King’s College London, De Crespigny Park, London, SE5 8AF, UK.; MRC London Institute of Medical Sciences, Hammersmith Hospital, London W12 0NN.; Institute of Clinical Sciences (ICS), Faculty of Medicine, Imperial College London, Du Cane Road, London W12 0NN.; Department of Psychiatry and Behavioral Sciences, Johns Hopkins Medical Institutions, Baltimore, MD, USA.; Russell H. Morgan Department of Radiology and Radiological Science, Johns Hopkins Medical Institutions, Baltimore, MD, USA.; University of Toronto, Department of Psychiatry, Toronto, Canada.; Department of Neuroimaging, Institute of Psychiatry, Psychology & Neuroscience, King’s College London, London, UK.

**Keywords:** positron emission tomography, psychosis, schizophrenia, translocator protein, microglia, immuneactivation, meta-analysis

## Abstract

**Background:** Accumulating evidence suggests that the immune system may be an important target for new treatment approaches in schizophrenia. Positron emission tomography (PET) and radioligands binding to the translocator protein (TSPO), which is expressed in glial cells in brain including immune cells, represents a potential method for patient stratification and treatment monitoring. This study examined if patients with first episode psychosis and schizophrenia had altered TSPO levels as compared to healthy control subjects.

**Methods:** PubMed was searched for studies comparing patients with psychosis to healthy controls using second-generation TSPO radioligands. The outcome measure was distribution volume (V_T_), an index of TSPO levels, in frontal cortex (FC), temporal cortex (TC) and hippocampus (HIP). Bayes factors (BF) were applied to examine the relative support for higher, lower or no-change of TSPO levels in patients as compared to healthy controls.

**Results:** Five studies, with 75 patients with first-episode psychosis or schizophrenia and 77 healthy control subjects were included. BF showed strong support for lower patient V_T_ relative to no-change (all BF>32) or relative to an increase (all BF>422) in all brain regions. From the posterior distributions, mean patient-control differences in standardized V_T_ values were −0.48 for FC (95% credible interval (CredInt)=-0.88 to −0.09), −0.47 for TC (CredInt=−0.87 to −0.07) and −0.63 for HIP (CredInt=−1.00 to −0.25).

**Discussion:** The observed reduction of TPSO in compared to control subjects in patients may correspond to altered function or lower density of brain immune cells. Future studies should focus on investigating the underlying biological mechanisms and their relevance for treatment.

## Introduction

Genetic, epidemiological and biomolecular data suggest that the immune system is involved in the pathophysiology of schizophrenia (1-3). When translating these findings into clinical trials, initial studies have shown positive effect of medication targeting the immune system when used as addon treatment to antipsychotics (4-6). To aid further development of this therapeutic approach, tools for directly assessing the status of the brain immune system are needed to allow for patient stratification and monitoring of treatment effects.

Using Positron Emission Tomography (PET), the localization and activation state of central nervous system (CNS) immune response modulators can be assessed with radioligands targeting the glial cell marker 18 kDa translocator protein (TSPO) (7-9). During the last decade, a handful of TSPO PET studies have been performed in patients with early-stage psychosis or manifest schizophrenia, showing inconclusive results. Early reports using the first-generation TSPO radioligand (R)-[^11^C]PK11195 showed increased binding in small patient groups (n=7 and n=10) (10, 11), albeit with outcome measures that show low accuracy and reliability (i.e. binding potential estimated from rate constants) (12-14). More recent studies in larger samples using the same radioligand, but without blood sampling for full quantification, did not replicate these findings (15-17). Concerns regarding the low signal to noise ratio of (R)-[^11^C]PK11195 sparked the development of a series of second-generation TSPO radioligands, showing much greater specific binding (18-21). These tools have subsequently been used to revisit the question of TSPO increases in psychosis (22-26). When employing gold standard outcome measures of binding in the absence of a reference region (distribution volume, V_T_, obtained using kinetic modeling with metabolite-corrected arterial plasma as input function), no increases have thus far been found in patients. In some cases, trend-level (24) or significantly lower TSPO levels (23) were shown.

All previous TSPO PET studies in psychosis have been performed with relatively small sample sizes. In addition, TSPO radioligands display a substantial within- and between-subject variability (12, 27), even after accounting for the TSPO rs6971 polymorphism which is known to affect radioligand binding in vivo (28-30). This has important implications for sensitivity and the power to detect differences between psychotic patients and controls. Indeed, the power to detect an expected significant medium-sized difference between diagnostic groups (at alpha=0.05) has ranged from 23% to 34% in previous designs (22-26). Medication status has also differed both between and within these studies. Since antipsychotics have been shown to dampen the immune response, this further limits the conclusions that can be drawn (31). Here, we sought to overcome these limitations and clarify the use of TSPO PET as a biomarker of immune dysfunction in schizophrenia. We conducted an individual participant data (IPD) meta-analysis of all TPSO PET studies performed in psychosis or schizophrenia using second-generation radioligands, where V_T_ was included as the outcome measure. The primary objective was to evaluate the hypotheses of 1) an increase or 2) a decrease or 3) no difference in V_T_ between patients and healthy control subjects. A secondary objective was to assess the effects of antipsychotic medication on TSPO binding.

## Materials and Methods

### PRISMA, pre-registration and code availability

This meta-analysis was conducted according to the Preferred Reporting Items for Systematic Reviews and Meta-analyses of Individual Participant Data (PRISMA-IPD) (32) and according to a study specific pre-registration protocol. The pre-registration protocol and all code used in this study can be found on the public repository https://github.com/pontusps/TSPO_psychosis.

### Selection Criteria and Search Strategy

We set out to obtain individual participant data from all PET studies that 1) used a second-generation TSPO radioligand, 2) reported distribution volume (V_T_) values in the CNS in patients with psychosis as compared to healthy controls (HC), and 3) reported TSPO affinity type of all participants. To our knowledge there are currently five published studies reporting such data, using the radioligands [^11^C]PBR28, [^18^F]FEPPA and [^11^C]DPA713 (22-26). In order to ascertain that no relevant studies were omitted from this meta-analysis, we performed a systematic literature search on PubMed. Only articles published after 2004 were included in the search, corresponding to the year when the first report on a second-generation TSPO radioligand was published (33). Search terms included, among others: “psychotic disorder”, “schizophrenia”, “positron emission tomography”, “translocator protein 18 kDa” and “peripheral benzodiazepine receptor” (for full list of search terms see Supplementary Information 1). All TSPO PET studies in psychosis or schizophrenia which were not included are listed in supplementary Table 1, along with a detailed explanation of the selection criteria. Corresponding authors of eligible studies were contacted via email and all agreed to contribute.

### Requested data

Requested IPD included V_T_ values from the Frontal Cortex (FC), Temporal Cortex (TC) and Hippocampus (HIP) regions of interest (ROIs), patient-control status, TSPO genotype, age, sex and medication status, Positive And Negative-Syndrome Scale in Schizophrenia (PANSS) scores (or equivalent) and duration of illness. These three ROIs were selected since four out of five included studies had reported V_T_ from all of them. Upon request, the corresponding author of the remaining study (22), provided unpublished IPD from all three ROIs to allow for consistent pooling. In order to account for range differences between different radioligands used across studies, we z-scored all ROI V_T_ values within each genotype group of each study.

### Quality control

The first author (PPS) examined the integrity of the obtained IPD datasets. The data was checked for outliers and inconsistencies to the published data (such as number of participants, means, ranges, and SDs of V_T_ and age), which were then resolved following discussion with the authors of the relevant study.

### Meta-analysis and statistics

The studies included in this meta-analysis recruited participants of two different TSPO affinity types (high-affinity binders, HABs; and mixed-affinity binders, MABs), used different radioligands, and applied different image analysis procedures. In order to estimate the difference in V_T_ between diagnostic groups (ΔV_T_) while taking this hierarchical structure into account, we constructed and compared four different Bayesian linear mixed effect (BLME) models of increasing complexity: M1) standardized ROI V_T_ was specified as dependent variable, diagnostic group as fixed effect, genotype and study as random effects with varying intercepts; M2) The same as M1 but with varying slopes of the random effect of genotype (i.e. allowing for differences in ΔV_T_ between HABs and MABs); M3) The same as M1 but with varying slopes of the random effect of study (i.e. allowing for differences in ΔV_T_ between studies); M4) The same as M1 but with varying slopes for both random effects (i.e. allowing for differences in ΔV_T_ between genotypes and studies). The model with the best fit to data, as determined by Widely Applicable Information Criterion (WAIC) and Leave-One-Out Cross-validation (LOOC) scores, was selected (34).

Following model selection, we first examined the hypothesis that patients have higher TSPO binding in the brain (H1). For each ROI we quantified the relative evidence of elevated TSPO expression in patients compared to the null-hypothesis of no change (H0). This was done using order-restricted Bayes Factor (BF) hypothesis testing (35-37) on ΔV_T_. BF quantifies the relative evidence, or support, for one hypothesis over another as a ratio of their average likelihoods. A BF > 10 is usually considered as strong evidence in favor of a hypothesis (and, consequently, BF < 0.1 translates into strong evidence of the opposite hypothesis) (35). We calculated BF_H1:H0_ to quantify the evidence in favor of an elevated ROI V_T_ signal, compared to no change, in patients. Secondly, we examined whether patients had lower levels of V_T_ in the ROI (H2). Again, this was done by employing an order-restricted BF test of decreased V_T_ in patients (BF_H2:H0_) over no change. Finally, we calculated the support for H2 over H1 (BF_H2:H1_), signaling how much more likely a decrease in patient TSPO binding is compared to an increase.

For each ROI, H1 and H2 were specified as half-Gaussian (normal) distributions centered on zero with a standard deviation of 0.5. Hence, in order to perform order-restricted hypothesis testing of patient-control differences, the priors over ROI ΔVτ were specified as half-Gaussians (SD=0.5) with a lower bound of zero for H1, and an upper bound of zero for H2. The Savage-Dickey Ratio method was then used to calculate BFs. The standard deviation was set a-priori to 0. 5 as this assigns high plausibility to ΔV_T_ values ranging from 0 to a medium-sized difference (38, 39). A medium-sized difference was considered a reasonable prediction, based on the precision of the outcome measure (27).

A robustness check of the effect of different prior widths on BF was performed by varying the SDs of the half-Gaussian distributions (SD = 0.2 and 0.8 – corresponding to an expected small and large effect size of ΔV_T_, respectively) when testing all hypotheses. For the prior on the SDs of the random effects, half-Cauchy distributions (with a scale of 0.707) were used. These weakly informative priors were chosen as the numbers of genotype groups (n=2) and studies (n=5) are small (40).

We also estimated the overall effect size of standardized V_T_ difference between patients and HC. This was done using model M3 with a non-truncated, weakly regularizing prior (Gaussian with a SD of 10) over the fixed effect. M3 was selected since it also allowed us to extract the study specific effects of ROI ΔV_T_ (random slopes), and the corresponding SD of these effects (τ). Using these we produced a “forest plot” of ROI ΔV_T_ and examined τ as a measure of study-heterogeneity, in line with the PRISMA-IPD guidelines.

For the secondary aim of analyzing medication effects on V_T_, we added an additional predictor, denoting medication status, to the best fitting BLME model. This predictor quantifies the additional effect of being medicated, after controlling for patient-control status. For each ROI, the prior distribution over the beta coefficient was a non-truncated Gaussian centered on zero with a SD of 10. The posterior of this predictor was then extracted together with its summary statistics (mean and 95% credible intervals (CredInt95%)) to examine the effect of medication.

We also examined the correlation between ROI V_T_ values and PANSS-Positive, PANSS-Negative scores as well as duration of illness (DOI) using linear effect modelling, allowing the correlations to vary between studies. All data was z-transformed within study (and within genotype for V_T_) and a uniform prior, ranging from −1 to 1, was specified for the beta coefficient.

In addition, frequentist equivalents of the best fitting model, showing p-values for patient-control differences in standardized V_T_ for each ROI are presented in Supplementary Table 2. The Hamiltonian Markov Chain Monte Carlo sampler STAN (41), and the R-packages brms (42) and lme4 (43) were used for the statistical modeling in this meta-analysis.

## Results

### Study selection and data collection

The PubMed search was performed on the 20th of February 2017 and resulted in thirteen research articles. The articles were read in full by two authors (PPS and SC). Both authors concluded independently that five studies (22-26) fulfilled the inclusion criteria for this meta-analysis (see PRISMA-flowchart in Supplementary Information 2). Each corresponding author provided anonymized individual participant V_T_ values from the frontal cortex (three studies (22-24)), dorsolateral prefrontal cortex (DLPFC) (two studies (25, 26)), temporal cortex (all studies) and hippocampus (all studies). For all subsequent analyses in this study, the V_T_ values from FC and DLPFC were considered to represent the same ROI.

### Characteristics of studies and quality control

Table 1 shows demographic information, medication status, PANSS (or equivalent), and duration of illness of all participants included in this meta-analysis. In total, IPD from 75 patients and 77 HC subjects were included in the statistical analysis. All patients in Kenk et al. (26), Bloomfield et al. (22) and all patients except two in Coughlin et al. (24) were on anti-psychotic treatment at the time of PET. Of the 19 patients who participated in Hafizi et al. (25), 5 patients were antipsychotic free with less than 4 weeks lifetime cumulative exposure, and 14 patients were antipsychotic naïve at the time of scanning. All patients in Collste et al. (23) were anti-psychotic naive. For all studies, exclusion criteria included clinically significant medical comorbidity and substance abuse. In two of the studies benzodiazepines were not allowed (22, 24), whereas in Collste et al. (23), and Kenk et al. (26) the results did not change when removing patients using benzodiazepines. Based on this information, as well as in vitro data showing effects of only high doses of diazepam on TSPO binding (44), we chose not to include this variable in our analysis. Figure 1 displays the individual participant ROI V_T_ values from the five studies included in this meta-analysis.

**Figure 1:**
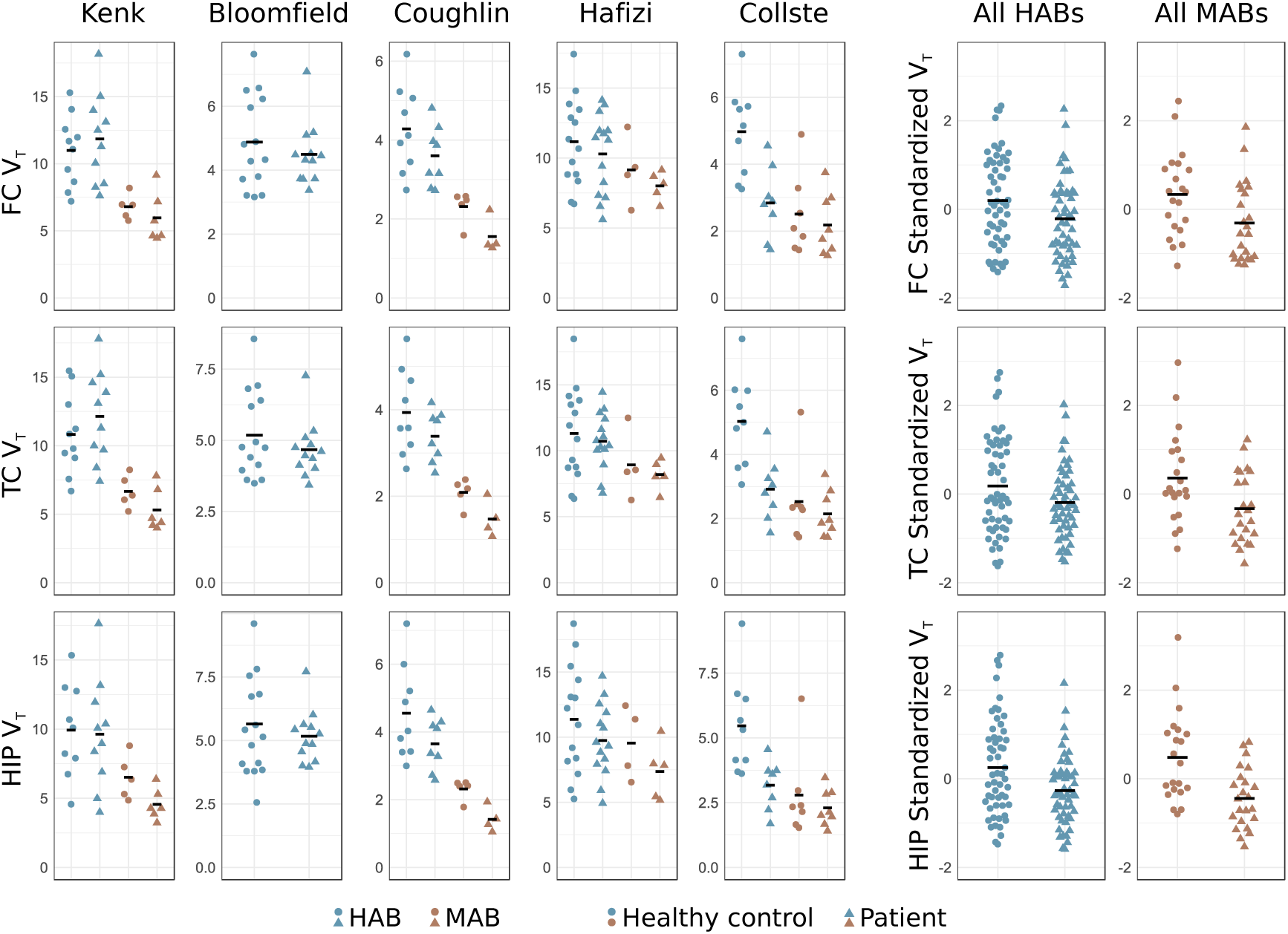
Individual participant raw data showing TSPO binding (estimated using V_T_) in patients with psychosis disorder and healthy controls, from all five included studies, from frontal cortex (FC), temporal cortex (TC) and hippocampus (HIP). The black bars denote the group means. For each region, subjects’ V_T_ values have been z-scored within study, and within genotype, in order to produce the pooled plots of all HABs and MABs. For this reason, HABs and MABs have the same mean (set to zero) in the right-hand panels.

**Table 1:** Descriptive characteristics of included data.

| Diagnostic group<sup>a</sup> | Schizophrenia /Other<sup>bc</sup> | Age Mean | Age SD | Count | HABs | MABs<sup>d</sup> | Males | Females | PANSS-T | PANSS-P<sup>e</sup> | PANSS-N | DOI Mean (Months) | Drugfree /Total |
| --- | --- | --- | --- | --- | --- | --- | --- | --- | --- | --- | --- | --- | --- |
| **Bloomfield et al.** |  |  |  |  |  |  |  |  |  |  |  |  |  |
| HC | - | 46.21 | 13.62 | 14 | 14 | 0 | 11 | 3 | - | - | - | - | - |
| pat | 12/0 | 47.00 | 9.31 | 12 | 12 | 0 | 9 | 3 | 63.7 (18.1) | 17.0 (6.1) | 14.1 (4.0) | 108.9 (46.7) | 0/12 |
| **Collste et al.** |  |  |  |  |  |  |  |  |  |  |  |  |  |
| HC | - | 26.38 | 8.44 | 16 | 9 | 7 | 7 | 9 | - | - | - | - | - |
| pat | 4/12 | 28.50 | 8.37 | 16 | 8 | 8 | 11 | 5 | 77.4 (18.3) | 20.3 (4.9) | 18.1 (7.0) | 7.9 (9.6) | 16/16 |
| **Coughlin et al.** |  |  |  |  |  |  |  |  |  |  |  |  |  |
| HC | - | 25.36 | 4.89 | 14 | 9 | 5 | 9 | 5 | - | - | - | - | - |
| pat | 12/0 | 24.33 | 3.28 | 12 | 8 | 4 | 9 | 3 | - | 13.8 (2.7) | 15.8 (4.6) | 25.0 (16.3) | 2/12 |
| **Hafizi et al.** |  |  |  |  |  |  |  |  |  |  |  |  |  |
| HC | - | 27.17 | 9.07 | 18 | 14 | 4 | 8 | 10 | - | - | - | - | - |
| pat | 15/4 | 27.53 | 6.78 | 19 | 14 | 5 | 12 | 7 | 68.6 (13.0) | 19.2 (3.8) | 16.1 (6.1) | 33.6 (40.1) | 19/19 |
| **Kenk et al.** |  |  |  |  |  |  |  |  |  |  |  |  |  |
| HC | - | 54.27 | 9.51 | 15 | 10 | 5 | 7 | 8 | - | - | - | - | - |
| pat | 16/0 | 42.50 | 14.03 | 16 | 10 | 6 | 10 | 6 | 70.2 (9.7) | 19.3 (2.2) | 18.6 (5.0) | 177.3 (105.7) | 0/16 |
| **All** |  |  |  |  |  |  |  |  |  |  |  |  |  |
| HC | - | 35.42 | 15.12 | 77 | 56 | 21 | 42 | 35 | - | - | - | - | - |
| pat | 59/16 | 33.88 | 12.57 | 75 | 52 | 23 | 51 | 24 | - | 18.2 (4.2) | 16.6 (5.5) | 72.1 (57.2) | 37/77 |

<sup>a</sup> The 14 HC subjects shared across the Kenk et al. and Hafizi et al. have been uniquely assigned to either one of the studies.
<sup>b</sup> Collste et al. other diagnoses: 7 schizophreniform disorder, 4 psychosis NOS, 1 brief psychosis.
<sup>c</sup> Hafizi et al. other diagnoses: 3 schizophreniform and 1 delusional disorder
<sup>d</sup> The 2 MAB patients were excluded from the hierarchical inferential analyses since z-scoring within genotype was not meaningful
<sup>e</sup> PANSS – P score converted from SAPS score, and PANSS – N score converted from SANS score, using van Erp et al., 2014

Healthy control subjects were recruited by flyers (Kenk et al. (26), Coughlin et al (24). and Hafizi et al. (25)), advertising in newspapers (Bloomfield et al. (22)), word of mouth (Coughlin et al. (24)) and advertising on internet (Kenk et al. (26), (25) and Collste et al. (23)) Exclusion criteria for all healthy controls included history of psychiatric disease or other clinically significant medical illness. Fourteen HC subjects from Kenk et al. (26) also served as controls in Hafizi et al. (25). Since different image analysis procedures were used in the two studies, it was not possible to employ a multiple membership model to account for this overlap. Instead, we assigned these 14 subjects to either the Kenk et al. (26) or the Hafizi et al. (25) data set, to make sure that data from the same subject was not used twice in the model. The assignment was performed prior to the inferential analyses, with the purpose of finding the best possible match between the diagnostic groups within both studies. In addition, one HC subject in Kenk et al. (26) had an outlier HIP V_T_ value (75.55), and a mismatch in MAB patient count was found in the Bloomfield et al. (22) data. These inconsistencies were resolved after consultation with the original authors. The final data set from Bloomfield et al. (22) contained two MAB patients, but no MAB HC. These two patients were excluded from the inferential analyses as standardization (z-scoring) was not meaningful.

The mean age of all patients was 33.88 (SD=12.57) and the mean age of healthy controls was 35.42 (SD=15.12). This corresponds to a negligible difference in age between diagnostic groups (Cohen’s d=0.11). Fisher’s exact test indicated some skewness in gender distribution between the patient and control groups (p=0.0504). In order to ascertain that any potential differences in ROI V_T_ values between diagnostic groups in the main analysis were not driven by gender differences, we included gender as a covariate, and executed an additional set of BLME models, using the same procedure as outlined in the methods.

### Model selection

Model M1 showed a slightly better fit, determined by WAIC and LOOC scores, compared to M2 and M3 (Table 2). We therefore used model M1 to obtain order-restricted posterior distributions of ROI ΔV_T_ and subsequently quantified evidence in favor of H0, H1 and H2.

**Table 2:**
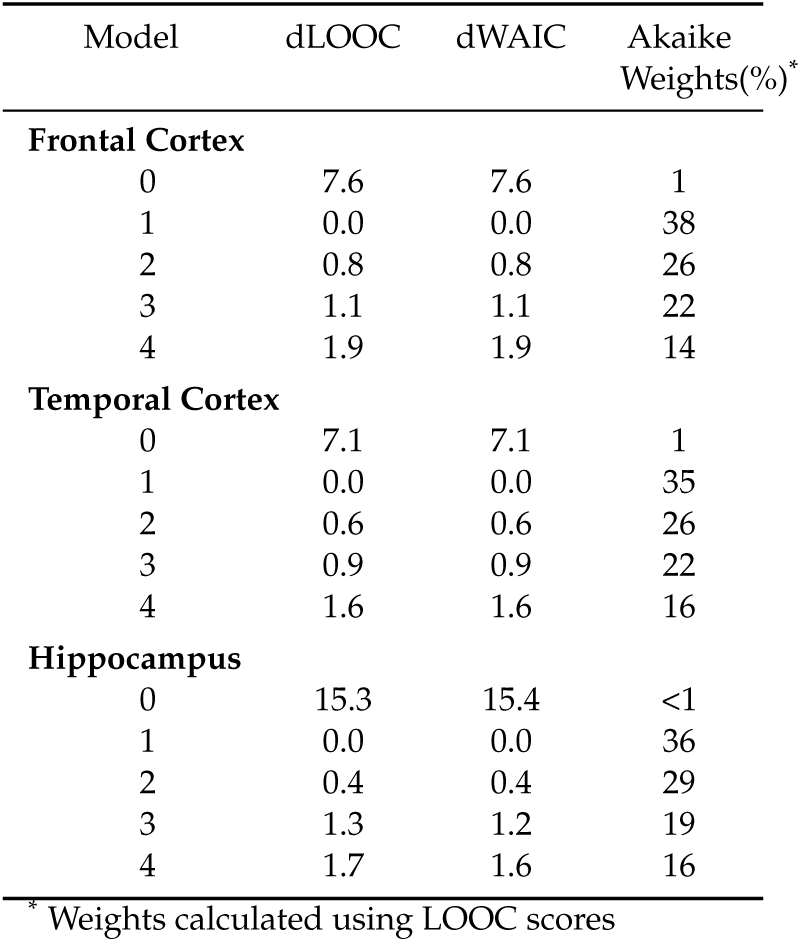
Model fits for four different Bayesian linear mixed effect models examining the difference in TSPO binding (estimated using V_T_) between patients with psychosis and healthy controls. A null model (0) without patient-control status as predictor is included as a baseline comparison. dLOOC is the distance to the best fitting model calculated using Leave-One-Out Cross-Validation; dWAIC is the distance to best fitting model calculated using the Widely Applicable Information Criteria. Lower dLOOC and dWAIC values indicate better model fit.

### Patient and control difference in V_T_ (primary aim)

BFH1:H0 in favor of elevated V_T_ in patients (H1) were 0.08 for FC, 0.08 for TC and 0.06 for hip-pocampus. This translates into strong support for the null-hypotheses of no change (H0) relative to an increase in patients. BF_H2:H0_ in favor of decreased V_T_ in patients (H2) were 32.5 for FC, 34.2 for TC and 1481.0 for hippocampus, compared to H0. This signifies very strong evidence for a lower V_T_ in patients. As a result, there was extremely strong support for H2 over H1 (BFH2:H1 FC: 422.9; TC: 440.6; hippocampus: 24524.0). Hence, a decreased V_T_ in psychosis patients, as compared to healthy controls, is over 422 times more likely than an increased V_T_, conditioned on the data and the models (see Table 3 and Supplementary Figure 1 for all computed BFs).

**Table 3:** Bayes factors of hypothesis testing of the difference in standardized brain TSPO binding (estimated using VT) between patients and controls, using the best fitting model (M1).

| Region* | H0:H1 | H1:H0 | H0:H2 | H2:H0 | H1:H2 | H2:H1 |
| --- | --- | --- | --- | --- | --- | --- |
| FC | 13.0 | 0.08 | 0.030 | 32.5 | 0.002 | 422.9 |
| TC | 12.9 | 0.08 | 0.030 | 34.2 | 0.002 | 440.6 |
| HIP | 16.6 | 0.06 | 0.001 | 1481.0 | <0.001 | 24524.0 |

\* FC = frontal cortex; TC = temporal cortex; HIP = hippocampus

When varying the widths (SD=0.2 and SD=0.8) of the Gaussian prior distribution on the fixed effect of differences between patients and controls, there was still strong support in favor of H2 for all ROIs (all BF_H2:H0_>15, see Supplementary Table 3). The addition of gender as a covariate did not change the qualitative inference for any of the ROIs (all BF_H2:H0_>16, see Supplementary Table 4).

### Estimation of effect sizes and study heterogeneity

For estimation of effect sizes and study heterogeneity, model M3, with an uninformative prior over ΔV_T_, was used. Figure 2 displays forest plots of the estimated patient-control difference in each study for each ROI. It also shows the posterior distributions of the standardized ΔV_T_ across all studies, together with summary statistics (mean and credible intervals). The mean of each ROI’s posterior distribution corresponds to a medium-sized difference in V_T_ between patients and controls.

**Figure 2:**
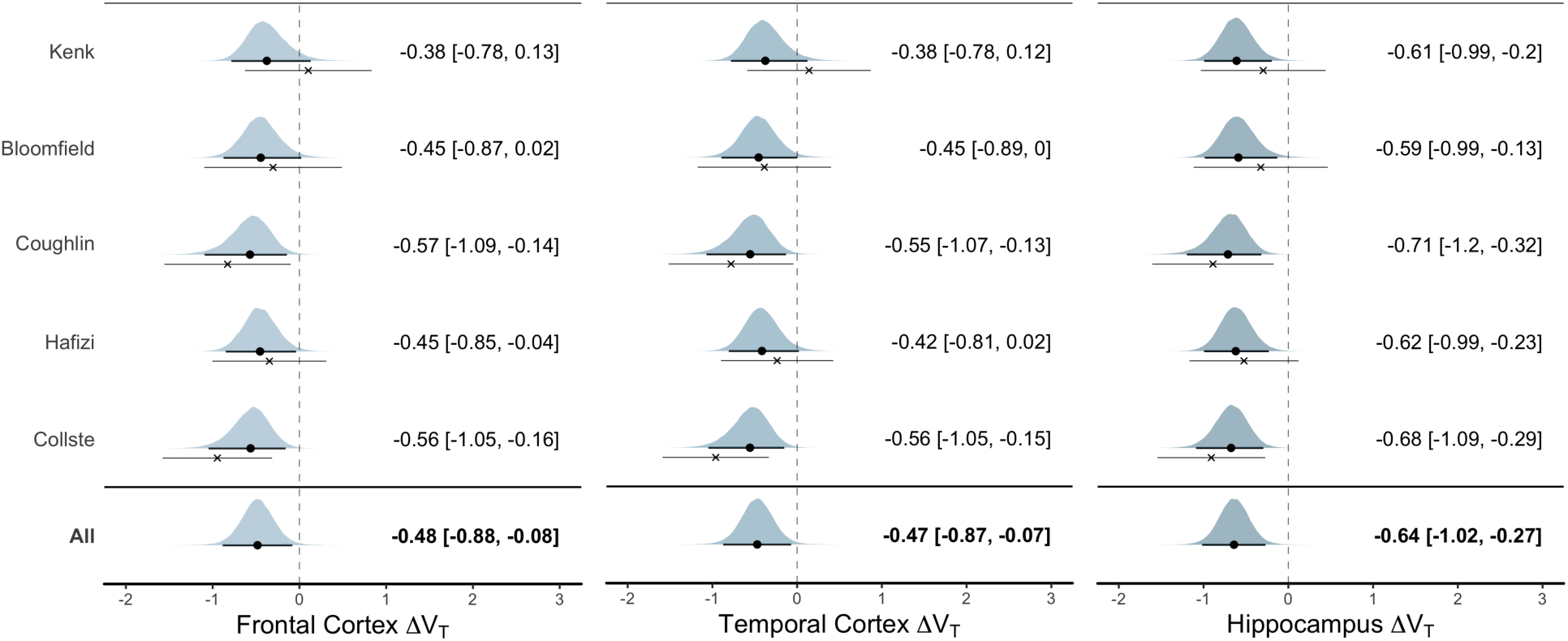
Standardized difference in TSPO binding (estimated using V_t_) between patients with psychosis disorder and healthy controls. The posterior distribution for each study-specific ΔV_T_ estimate (random slopes) from the linear mixed model are presented. The black circle denotes the posterior mean, and the thick line the 95% credible interval, which are also presented in text next to the plots. The cross denotes the patient-control mean difference in raw data (together with its 95% CI), without performing linear mixed effects modeling. Hence, the difference between the dot and the cross displays the model shrinkage towards the mean. The overall ΔV_T_ estimate shows a clear decrease in patients’ TSPO binding, compared to healthy controls.

For all ROIs, the SDs of the random slopes of studies (τ) were very small (posterior modes<0.04; posterior means<0.22,) and I^2^<15%, signifying low study heterogeneity in ΔV_T_ differences (see Supplementary Figure 2).

### Effect of medication (secondary aim)

We examined the effect of medication on V_T_, by adding medication-status as an additional predictor to model M1. For all ROIs, the models showed little to no evidence of a medication effect, allocating as much probability to an increase as to a decrease. The mean of the posterior over the change in standardized V_T_ due to medication was 0.009 for FC (CredInt95% −0.384 to 0.401), −0.013 for TC (CredInt95% −0.407 to 0.381) and −0.040 for HIP (CredInt95% −0.423 to 0.343), see Figure 3. Thus, no support was found for a difference between drug-free and medicated patients.

**Figure 3:**
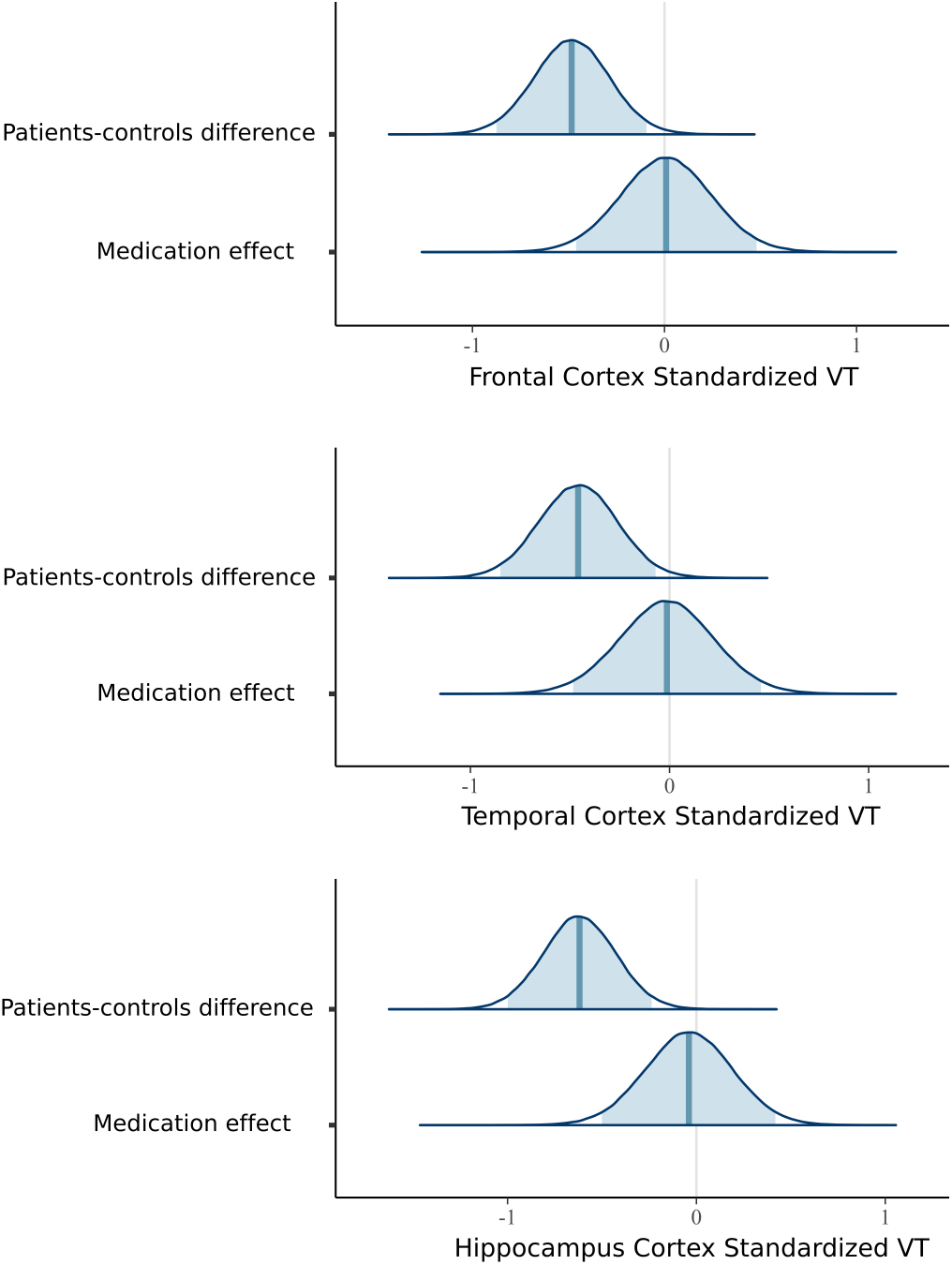
Posterior distributions over the differences in standardized brain TSPO binding (estimated using V_T_) values between patients and controls, and the additional effect of medication status (being medicated with anti-psychotics or not at the time of PET). The posterior distributions of medication effect are centred on zero and suggest that anti-psychotic treatment does not affect brain V_T_, after differences between psychosis patients and controls have been accounted for.

### Correlation to PANSS and DOI

There was little to no evidence for a correlation between regional V_T_ and PANSS-Positive scores (posterior means=0.02 to 0.06, p=0.61 to 0.86) as well as between regional V_T_ and PANSS-Negative scores (posterior means=0.02 to 0.05; p=0.71 to 0.88). See Supplementary Figure 4 and Supplementary Table 5 for full reporting of V_t_ and PANSS correlations. There was little to no evidence for a correlation between regional V_t_ and DOI (posterior means = −0.11 to −0.07; p = 0.28 to 0.52). See Supplementary Figure 5 and Supplementary Table 6 for full reporting of V_t_ and DOI correlations.

## Discussion

The main finding of this IPD meta-analysis was a reduction in binding of the glial cell marker TSPO in schizophrenia and first-episode psychosis compared to healthy control subjects. Using Bayesian linear-mixed-effect modeling, we observed very strong evidence of lower TSPO binding, measured using V_t_, in the FC, TC and hippocampus, contrary to the hypothesis of elevated TSPO in patients. As such, this study constitutes the most conclusive in vivo investigation of TSPO in psychosis to date.

Antipsychotic medication has been shown to attenuate blood cytokine levels in patients (31) as well as inhibit immune cell activity in vitro (45). Although the effect on TSPO expression in animals is less conclusive (46), these observations suggest that TSPO binding could be lower in medicated compared to unmedicated subjects. However, our secondary analysis of the effect of medication status yielded no evidence for such a difference in TSPO binding between drug-free and medicated patients. This indicates that the observed reduction is not an effect of exposure to antipsychotic treatment.

A wealth of data have demonstrated increases in pro-inflammatory markers, such as cytokines, in CSF and plasma in patients across disease stages of schizophrenia (3, 47). In the brain, these signaling molecules are mainly released by microglia and astrocytes, which have key roles in the immune response (9). Therefore, increases in numbers or activity of these cells in schizophrenia has been hypothesized (48,49). In post-mortem studies, increases in brain glial cell markers such as HLA-DR and CD11b have been observed in patients, although results have been mixed (50-52). With regard to astrocyte markers, there is no evidence of any overall differences between patients and controls (51, 52). In the case of TSPO, which is expressed in microglia and astrocytes (8, 9, 53), autoradiographic studies have reported both higher (28) and lower (54) binding in patients as compared to healthy controls. Important caveats, when interpreting these studies, are that the age of patients and control subjects is generally high, and in patients the cause of death is often suicide (52). A recent translational study examined TSPO in an infection-mediated animal model of schizophrenia. Increases in pro-inflammatory cytokines were found in brain regions which also showed reductions in TSPO levels as measured using immunohistochemistry (55), an observation that paralleled TSPO PET and CSF data in patients (24). Importantly, microglia and astrocytes have been found to exist in both pro- and anti-inflammatory states (56, 57), which cannot be differentiated by TSPO. Indeed, very recent in vitro data suggest that M1 (pro-inflammatory) macrophages may show reduced TSPO expression (58). The above discussed literature, together with the results of our study, challenges the utility of TSPO as an exclusively pro-inflammatory marker in schizophrenia. Lower TSPO binding could indicate a compensatory mechanism to a pro-inflammatory signal (55, 59), or altered function of glial cells such as abnormal energy utilization (60). Since stimulation of TSPO has shown to attenuate microglial activation in response to neuroinflammatory challenges (61-63), lower TSPO in psychosis could also indicate an inherent weaker anti-inflammatory response. These hypotheses all need to be addressed in future studies.

Since there is no brain region devoid of TSPO expression (64, 65), metabolite-corrected arterial plasma measurements of radioligand concentration are necessary for accurate in vivo quantification of binding. In order to overcome variability that may be associated with the arterial measurements (27, 66), relative measures of binding, such as distribution volume ratios (DVRs), have been proposed (22). Out of the studies included in this meta-analysis, one reported a significant elevation of DVR in schizophrenia patients and people at clinical risk for psychosis (22) whereas three studies showed no difference in schizophrenia (23–25). More recently, no increase in DVR was found in high risk individuals (67). We chose not to include DVR in our analysis. The interpretation of patient-control differences obtained by dividing binding in a target region with that of a reference region are complicated by the possibility that there could be alterations in specific binding in the reference region as well. In addition, the reliability of DVR for TSPO radioligands has been found to be low (68). Given the lack of a true reference region, V_T_ is the most suitable outcome for TSPO quantification, but it is not without drawbacks. Apart from glial specific binding (Vs), the signal includes non-specific and free radioligand (V_ND_). A general caveat with all studies performed using V_T_ is that, without blocking data (69), it cannot be excluded that V_ND_ differs between groups. However, for TSPO imaging, this has yet to be shown for any disorder or condition. Also, TSPO binding is expressed in perivascular and endothelial cells (55, 70), and under certain conditions also neurons (71). Further research is needed to evaluate the contribution of these components to observed decrease in V_T_ in schizophrenia. Finally, although arterial measurement currently represents the most accurate estimate available of radioligand delivery to brain, it cannot be fully excluded that unknown factors affecting the accuracy of these measures could be unequally distributed between groups. Of the included studies, only Bloomfield et al. (22), reported changes in input function (increases, as shown in an ensuing publication (72)), whereas Collste et al. (23) and Hafizi et al. (25) found no group differences.

In this IPD meta-analysis, the hierarchal statistical models allowed us to investigate the difference in TSPO binding between patients with psychosis and healthy controls across five different studies. By including only studies employing second-generation radiotracers, and reporting the standard outcome measure V_T_, the analysis fulfils the pre-condition of meta-analytical models that outcomes should stem from the same underlying distribution of effects. Synthesizing data in this way, we were able to overcome the critical limitation of small sample sizes in the individual reports. Despite this, the total number of included subjects did not allow for investigations of specific subgroups, such as different disease stages.

## Conclusions

The present study shows that TSPO binding is reduced across several brain regions in patients with first-episode psychosis and schizophrenia, suggesting an altered function, or reduced density of immune and glial cells. Further work is needed to assess the exact biological meaning of these changes, using both clinical and translational studies.

## Acknowledgement

We thank Yong Du, Ph.D., and Sina Hafizi, M.D., Ph.D., for helpful comments on the manuscript. The pre-registration study- and analysis protocol together with all analysis code can be found at https://github.com/pontusps/TSPO_psychosis.

Dr Cervenka reports funds from the Swedish Research Council (523-2014-3467) and Stockholm County Council. Dr Collste reports funding from PRIMA Barn- och Vuxenpsykiatri AB. Dr Howes reports funds from Research Council-UK (no. MC-A656-5QD30), Maudsley Charity (no. 666), Brain and Behavior Research Foundation, and Wellcome Trust (no. 094849/Z/10/Z), and the National Institute for Health Research (NIHR) Biomedical Research Centre at South London and Maudsley NHS Foundation Trust and King’s College London. Dr Ashok reports funding from the Medical Research Council and King’s College London. Dr Pomper reports funding from US DoD (GW130098). Dr Mizrahi reports funding from the National Institutes of Health (NIH) R01 grant MH100043. The funders had no role in the design and conduct of the study; collection, management, analysis, and interpretation of the data; preparation, review, or approval of the manuscript; and decision to submit the manuscript for publication.

## Author contributions

Plavén-Sigray and Cervenka conceived of the study, design the study, wrote the study protocol and supervised the study. Plavén-Sigray and Cervenka carried out the literature search. Plavén-Sigray, Collste, Pompers, Coughlin, Wang, Mizrahi, Rusjan, Howes, Veronese, Ashok and Cervenka aided in the acquisition and quality control of data. Plavén-Sigray and Matheson performed the statistical analyses. Plavén-Sigray and Cervenka drafted the manuscript. All authors revised the manuscript for intellectual content and approved of the final version.

## References

1 Arias I, Sorlozano A, Villegas E, de Dios Luna J, McKenney K, Cervilla J et al. (2012): Infectious agents associated with schizophrenia: a meta-analysis. Schizophrenia research. 136: 128–136.

2 Ripke S, Neale BM, Pers TH, Julià A, Kahn RS, Kalaydjieva L et al. (2014): Biological insights from 108 schizophrenia-associated genetic loci| NOVA. The University of Newcastle’s Dig-ital Repository.

3 Upthegrove R, Manzanares-Teson N, Barnes NM (2014): Cytokine function in medication-naive first episode psychosis: a systematic review and meta-analysis. Schizophrenia research. 155: 101–108.

4 Akhondzadeh S, Tabatabaee M, Amini H, Abhari SAA, Abbasi SH, Behnam B (2007): Celecoxib as adjunctive therapy in schizophrenia: a double-blind, randomized and placebo-controlled trial. Schizophrenia research. 90: 179–185.

5 Müller N, Krause D, Dehning S, Musil R, Schennach-Wolff R, Obermeier M et al. (2010): Celecoxib treatment in an early stage of schizophrenia: results of a randomized, double-blind, placebo-controlled trial of celecoxib augmentation of amisulpride treatment. Schizophrenia research. 121: 118–124.

6 Rapaport MH, Delrahim KK, Bresee CJ, Maddux RE, Ahmadpour O, Dolnak D (2005): Celecoxib augmentation of continuously ill patients with schizophrenia. Biological psychiatry. 57: 1594–1596.

7 Ory D, Planas A, Dresselaers T, Gsell W, Postnov A, Celen S et al. (2015): PET imaging of TSPO in a rat model of local neuroinflammation induced by intracerebral injection of lipopolysaccharide. Nuclear medicine and biology. 42: 753–761.

8 Toth M, Little P, Arnberg F, Mulder J, Halldin C, Ha J, Holmin S (2015): Acute neuroinflammation in a clinically relevant focal cortical ischemic stroke model in rat : longitudinal positron emission tomography and immunofluorescent tracking. Brain Struct Funct. doi: 10.1007/s00429-014-0970-y.

9 Venneti S, Lopresti BJ, Wiley CA (2013): Molecular imaging of microglia/macrophages in the brain. Glia. 61: 10–23.

10 Doorduin J, De Vries EFJ, Willemsen ATM, De Groot JC, Dierckx RA, Klein HC (2009): Neuroinflammation in schizophrenia-related psychosis: a PET study. Journal of Nuclear Medicine. 50: 1801–1807.

11 Van Berckel BN, Bossong MG, Boellaard R, Kloet R, Schuitemaker A, Caspers E et al. (2008): Microglia activation in recent-onset schizophrenia: a quantitative (R)-[11 C] PK11195 positron emission tomography study. Biological psychiatry. 64: 820–822.

12 Jučaite A, Cselényi Z, Arvidsson A, Åhlberg G, Julin P, Varnäs K et al. (2012): Kinetic analysis and test-retest variability of the radioligand [11C](R)-PK11195 binding to TSPO in the human brain - a PET study in control subjects. EJNMMI research. 2: 1.

13 Slifstein M, Laruelle M (2001): Models and methods for derivation of in vivo neuroreceptor parameters with PET andSPECT reversible radiotracers. Nuclear Medicine and Biology. 28: 595–608.

14 Varnäs K, Varrone A, Farde L (2013): Modeling of PET data in CNS drug discovery and development. Journal of pharmacokinetics and pharmacodynamics. 40: 267–279.

15 Holmes SE, Hinz R, Drake RJ, Gregory CJ, Conen S, Matthews JC et al. (2016): In vivo imaging of brain microglial activity in antipsychotic-free and medicated schizophrenia: a [11C](R)-PK11195 positron emission tomography study. Molecular psychiatry. 21: 1672–1679.

16 Van Der Doef TF, De Witte LD, Sutterland AL, Jobse E, Yaqub M, Boellaard R et al. (2016): In vivo (R)-[11C]PK11195 PET imaging of 18kDa translocator protein in recent onset psychosis. NPJ schizophrenia. 2: 16031.

17 Di Biase MA, Zalesky A, O’keefe G, Laskaris L, Baune BT, Weickert CS et al. (2017): PET imaging of putative microglial activation in individuals at ultra-high risk for psychosis, recently diagnosed and chronically ill with schizophrenia. Translational psychiatry. 7: e1225.

18 Fujita M, Kobayashi M, Ikawa M, Gunn RN, Rabiner EA, Owen DR et al. (2017): Com-parison of four 11C-labeled PET ligands to quantify translocator protein 18 kDa (TSPO) in human brain: (R)-PK11195, PBR28, DPA-713, and ER176—based on recent publications that measured specific-to-non-displaceable ratios. EJNMMI Research. 7: 84.

19 Kobayashi M, Jiang T, Telu S, Zoghbi SS, Gunn RN, Rabiner EA et al. (2017): 11C-DPA-713 has much greater specific binding to translocator protein 18 kDa (TSPO) in human brain than 11C-(R)-PK11195. Journal of Cerebral Blood Flow & Metabolism. 0271678X17699223.

20 Kreisl WC, Fujita M, Fujimura Y, Kimura N, Jenko KJ, Kannan P et al. (2010): Comparison of [11C]-(R)-PK 11195 and [11C]PBR28, two radioligands for translocator protein (18 kDa) in human and monkey: Implications for positron emission tomographic imaging of this inflammation biomarker. Neuroimage. 49: 2924–2932.

21 Wilson AA, Garcia A, Parkes J, McCormick P, Stephenson KA, Houle S, Vasdev N (2008): Radiosynthesis and initial evaluation of [18F]-FEPPA for PET imaging of peripheral benzodiazepine receptors. Nuclear Medicine and Biology. 35: 305–314.

22 Bloomfield PS, Selvaraj S, Veronese M, Rizzo G, Bertoldo A, Owen DR et al. (2015): Microglial activity in people at ultra high risk of psychosis and in schizophrenia: an [11C] PBR28 PET brain imaging study. American Journal of Psychiatry.

23 Collste K, Plavén-Sigray P, Fatouros-Bergman H, Victorsson P, Schain M, Forsberg A et al. (2017): Lower levels of the glial cell marker TSPO in drug-naive first-episode psychosis patients as measured using PET and [11C]PBR28. Molecular Psychiatry. 22: 850–856.

24 Coughlin JM, Wang Y, Ambinder EB, Ward RE, Minn I, Vranesic M et al. (2016): In vivo markers of in fl ammatory response in recent-onset schizophrenia: a combined study using [11 C] DPA-713 PET and analysis of CSF and plasma. 1–8.

25 Hafizi S, Tseng HH, Rao N, Selvanathan T, Kenk M, Bazinet RP et al. (2017): Imaging microglial activation in untreated first-episode psychosis: A PET study with [18F]FEPPA. American Journal of Psychiatry. 174: 118–124.

26 Kenk M, Selvanathan T, Rao N, Suridjan I, Rusjan P, Remington G et al. (2015): Imaging neuroinflammation in gray and white matter in schizophrenia: an in-vivo PET study with [18F]-FEPPA. Schizophrenia bulletin. 41: 85–93.

27 Collste K, Forsberg A, Varrone A, Amini N, Aeinehband S, Yakushev I et al. (2016): Test-retest reproducibility of [11C]PBR28 binding to TSPO in healthy control subjects. European Journal of Nuclear Medicine and Molecular Imaging. 43: 173–183.

28 Kreisl WC, Jenko KJ, Hines CS, Lyoo CH, Corona W, Morse CL et al. (2013): A genetic polymorphism for translocator protein 18 kDa affects both in vitro and in vivo radioligand binding in human brain to this putative biomarker of neuroinflammation. Journal of Cerebral Blood Flow & Metabolism. 33: 53–58.

29 Owen DR, Yeo AJ, Gunn RN, Song K, Wadsworth G, Lewis A et al. (2012): An 18-kDa translocator protein (TSPO) polymorphism explains differences in binding affinity of the PET radioligand PBR28. Journal of Cerebral Blood Flow & Metabolism. 32: 1–5.

30 Owen DRJ, Gunn RN, Rabiner EA, Bennacef I, Fujita M, Kreisl WC et al. (2011): Mixed-affinity binding in humans with 18-kDa translocator protein ligands. Journal of Nuclear Medicine. 52: 24–32.

31 Drzyzga Ł, Obuchowicz E, Marcinowska A, Herman ZS (2006): Cytokines in schizophrenia and the effects of antipsychotic drugs. Brain, behavior, and immunity. 20: 532–545.

32 Stewart LA, Clarke M, Rovers M, Riley RD, Simmonds M, Stewart G, Tierney JF (2015): Preferred reporting items for a systematic review and meta-analysis of individual participant data: the PRISMA-IPD statement. Jama. 313: 1657–1665.

33 Maeda J, Suhara T, Zhang M, Okauchi T, Yasuno F, Ikoma Y et al. (2004): Novel peripheral benzodiazepine receptor ligand [11C]DAA1106 for PET: an imaging tool for glial cells in the brain. Synapse. 52: 283–291.

34 Vehtari A, Gelman A, Gabry J (2016): Practical Bayesian model evaluation using leave-one-out cross-validation and WAIC. Statistics and Computing. 1–20.

35 Jeffreys H (1961): Theory of probability, 3rd ed. Oxford: Oxford University Press.

36 Kass RE, Raftery AE (1995): Bayes factors. Journal of the american statistical association. 90: 773–795.

37 Morey RD, Wagenmakers E-J (2014): Simple relation between Bayesian order-restricted and point-null hypothesis tests. Statistics & Probability Letters. 92: 121–124.

38 Cohen J (1998): Statistical power analysis for the behavioral sciences. 2nd ed. Hillsdale, NJ: Erbaum.

39 Dienes Z (2014): Using Bayes to get the most out of non-significant results. Frontiers in psychology. 5: 781.

40 Gelman A (2006): Prior distributions for variance parameters in hierarchical models (comment on article by Browne and Draper). Bayesian analysis. 1: 515–534.

41 Carpenter B, Gelman A, Hoffman M, Lee D, Goodrich B, Betancourt M et al. (2016): Stan: A probabilistic programming language. Journal of Statistical Software. 20.

42 Buerkner P-C (2016): brms: An R package for Bayesian multilevel models using Stan. Journal of Statistical Software. 80: 1–28.

43 Bates D, Mächler M, Bolker B, Walker S (2014): Fitting linear mixed-effects models using lme4. arXiv preprint arXiv:14065823.

44 Kalk NJ, Owen DR, Tyacke RJ, Reynolds R, Rabiner EA, Lingford-hughes AR, Parker CA (2013): Are prescribed benzodiazepines likely to affect the availability of the 18 kda translocator protein (TSPO) in PET studies? Synapse. 67: 909–912.

45 Monji A, Kato TA, Mizoguchi Y, Horikawa H, Seki Y, Kasai M et al. (2013): Neuroinflammation in schizophrenia especially focused on the role of microglia. Progress in Neuro-Psychopharmacology and Biological Psychiatry. 42: 115–121.

46 Danovich L, Veenman L, Leschiner S, Lahav M, Shuster V, Weizman A, Gavish M (2008): The influence of clozapine treatment and other antipsychotics on the 18 kDa translocator protein, formerly named the peripheral-type benzodiazepine receptor, and steroid production. European Neuropsychopharmacology. 18: 24–33.

47 Miller BJ, Buckley P, Seabolt W, Mellor A, Kirkpatrick B (2011): Meta-analysis of cytokine alterations in schizophrenia: clinical status and antipsychotic effects. Biological psychiatry. 70: 663–671.

48 Howes OD, McCutcheon R (2017): Inflammation and the neural diathesis-stress hypothesis of schizophrenia: a reconceptualization. Translational psychiatry. 7: e1024.

49 Mizrahi R (2015): Social stress and psychosis risk: common neurochemical substrates? Neuropsychopharmacology.

50 Laskaris LE, Di Biase MA, Everall I, Chana G, Christopoulos A, Skafidas E et al. (2016): Microglial activation and progressive brain changes in schizophrenia. British journal ofpharmacology. 173: 666–680.

51 Trépanier MO, Hopperton KE, Mizrahi R, Mechawar N, Bazinet RP (2016): Postmortem evidence of cerebral inflammation in schizophrenia: a systematic review. Molecular psychiatry.

52 Kesteren C van, Gremmels H, Witte LD de, Hol EM, Van Gool AR, Falkai PG et al. (2017): Immune involvement in the pathogenesis of schizophrenia: a meta-analysis on postmortem brain studies. Translational Psychiatry. 7: e1075.

53 Lavisse S, Guillermier M, Hérard A-S, Petit F, Delahaye M, Van Camp N et al. (2012): Reactive astrocytes overexpress TSPO and are detected by TSPO positron emission tomography imaging. The Journal of Neuroscience. 32: 10809–10818.

54 Kurumaji A, Wakai T, Toru M (1997): Decreases in peripheral-type benzodiazepine receptors in postmortem brains of chronic schizophrenics. Journal of neural transmission. 104: 1361–1370.

55 Notter T, Coughlin JM, Gschwind T, Weber-Stadlbauer U, Wang Y, Kassiou M et al. (2017): Translational evaluation of translocator protein as a marker of neuroinflammation in schizophrenia. Molecular Psychiatry.

56 Liddelow SA, Guttenplan KA, Clarke LE, Bennett FC, Bohlen CJ, Schirmer L et al. (2017): Neurotoxic reactive astrocytes are induced by activated microglia. Nature.

57 Prinz M, Priller J (2014): Microglia and brain macrophages in the molecular age: from origin to neuropsychiatric disease. Nature Reviews Neuroscience. 15: 300–312.

58 Narayan N, Mandhair H, Smyth E, Dakin SG, Kiriakidis S, Wells L et al. (2017): The macrophage marker translocator protein (TSPO) is down-regulated on pro-inflammatory ‘M1’ human macrophages. PLOS ONE. 12: e0185767.

59 Forsberg A, Cervenka S, Jonsson Fagerlund M, Rasmussen LS, Zetterberg H, Erlandsson Harris H et al. (2017): The immune response of the human brain to abdominal surgery. Annals of Neurology.

60 Banati RB, Middleton RJ, Chan R, Hatty CR, Kam WW-Y, Quin C et al. (2014): Positron emission tomography and functional characterization of a complete PBR/TSPO knockout. Nature communications. 5.

61 Ryu JK, Choi HB, McLarnon JG (2005): Peripheral benzodiazepine receptor ligand PK11195 reduces microglial activation and neuronal death in quinolinic acid-injected rat striatum. Neurobi-ology of disease. 20: 550–561.

62 Leaver KR, Reynolds A, Bodard S, Guilloteau D, Chalon S, Kassiou M (2012): Effects of translocator protein (18 kDa) ligands on microglial activation and neuronal death in the quinolinic-acid-injected rat striatum. ACS chemical neuroscience. 3: 114–119.

63 Wang W, Zhang L, Zhang X, Xue R, Li L, Zhao W et al. (2016): Lentiviral-mediated overexpression of the 18 kDa translocator protein (TSPO) in the hippocampal dentate gyrus ameliorates LPS-induced cognitive impairment in mice. Frontiers in pharmacology. 7.

64 Doble A, Malgouris C, Daniel M, Daniel N, Imbault F, Basbaum A et al. (1987): Labelling of peripheral-type benzodiazepine binding sites in human brain with [3H] PK 11195: anatomical and subcellular distribution. Brain research bulletin. 18: 49–61.

65 Owen DR, Guo Q, Kalk NJ, Colasanti A, Kalogiannopoulou D, Dimber R et al. (2014): Determination of [11C]PBR28 binding potential in vivo: a first human TSPO blocking study. Journal of Cerebral Blood Flow & Metabolism. 34: 989–994.

66 Park E, Gallezot J-D, Delgadillo A, Liu S, Planeta B, Lin S-F et al. (2015): 11C-PBR28 imaging in multiple sclerosis patients and healthy controls: test-retest reproducibility and focal visualization of active white matter areas. European journal of nuclear medicine and molecular imaging. 42: 1081–1092.

67 Hafizi S, Da Silva T, Gerritsen C, Kiang M, Bagby RM, Prce I et al. (2017): Imaging Microglial Activation in Individuals at Clinical High Risk for Psychosis: an In Vivo PET Study with [(18) F] FEPPA. Neuropsychopharmacology: official publication of the American College of Neuropsy-chopharmacology.

68 Matheson G, Plavén-Sigray P, Forsberg A, Varrone A, Farde L, Cervenka S (2017): Assessment of simplified ratio-based approaches for quantification of PET [<sup>11</sup>C]PBR28 data. EJNMMI Research. 7. doi: 10.1186/s13550-017-0304-1.

69 Cunningham VJ, Rabiner EA, Slifstein M, Laruelle M, Gunn RN (2010): Measuring drug occupancy in the absence of a reference region: the Lassen plot re-visited. J Cereb Blood Flow Metab. 30. doi: 10.1038/jcbfm.2009.190.

70 Veronese M, Marques T, Bloomfield PS, Rizzo G, Singh N, Jones D et al. (n.d.): Kinetic modelling of [11C]PBR28 for 18kDa Translocator Protein PET data: a validation study of vascular modelling in the brain using XBD173 and tissue analysis. Journal of Cerebral Blood Flow and Metabolism.

71 Zhang M, Liu J, Zhou M-M, Wu H, Hou Y, Li Y-F et al. (2016): Elevated Neurosteroids in the Lateral Thalamus Relieve Neuropathic Pain in Rats with Spared Nerve Injury. Neuroscience bulletin. 32: 311–322.

72 Bloomfield PS, Howes OD, Turkheimer F, Selvaraj S, Veronese M (2016): Response to Narendran and Frankle: the interpretation of PET microglial imaging in schizophrenia. American Journal of Psychiatry. 173: 537–538.

